# The fermented cabbage metabolome and its protection against cytokine-induced intestinal barrier disruption of Caco-2 monolayers

**DOI:** 10.1101/2024.11.15.623851

**Authors:** Lei Wei, Maria L Marco

## Abstract

Fermented vegetables, such as fermented cabbage (sauerkraut), have garnered growing interest for their associations with a myriad of health benefits. However, the mechanistic details underlying the outcomes of consuming these foods require further investigation. This study examined the capacity of soluble metabolites in laboratory-scale and commercial fermented cabbage to protect against disruption of polarized Caco-2 monolayers by IFN-γ and TNF-α. Laboratory-scale ferments (LSF) were prepared with and without the addition of *Lactiplantibacillus plantarum* NCIMB8826R (LP8826R) and sampled after 7- and 14-days of incubation. Trans-epithelial electrical resistance (TER) and paracellular permeability to fluorescein isothiocyanate-dextran (FITC) revealed that fermented cabbage, but not raw cabbage or brine, protected against cytokine-induced damage to the Caco-2 monolayers. Barrier-protective effects occurred despite increased IL-8 production following cytokine exposure. Metabolomic analyses performed using gas and liquid chromatography resulted in the identification of 149 and 333 metabolites, respectively. Significant differences were found between raw and fermented cabbage. LSF metabolomes changed over time and the profiles of LSF with LP8826R best resembled the commercial product. Overall, fermentation resulted in lower carbohydrate and increased lactic acid, lipid, amino acid derivative (including D-phenyl-lactate (D-PLA), indole-3-lactate (ILA), and γ-aminobutyric acid (GABA)), and phenolic compound concentrations. Lactate, D-PLA, and ILA tested individually and combined only partially protected against cytokine-induced TER reductions and increases in paracellular permeability of Caco-2 monolayers. The findings show that intestinal barrier-protective compounds are consistently enriched during cabbage fermentations, irrespective of scale or microbial additions, that may contribute to the health-promoting potential of these foods.

**Importance:** Fermented vegetables are increasingly associated with health benefits. However, the importance of microbial transformations to foods during the fermentation process remain to be determined. We found that the metabolites in spontaneously fermented cabbage protected polarized intestinal epithelial cells against damage induced by proinflammatory cytokines. Cabbage fermentations resulted in consistent metabolome profiles enriched in bioactive compounds known to be made by beneficial members of the human gut microbiome, including D-phenyl-lactate (D-PLA) and indole-3-lactate (ILA). The metabolomes were distinct from raw cabbage and further differentiated based on sampling time, addition of an exogenous *Lactiplantibacillus plantarum* strain, and source. Because only partial protection against intestinal barrier disruption was found when individual metabolites (D-PLA, ILA, lactate) were applied, the findings indicate that the complex mixture of metabolites in a cabbage fermentation offer advantages over single metabolites to benefit intestinal health.

## Introduction

Fermented vegetables have been staples of the human diet for thousands of years (Medina-Pradas et al., 2017). Besides imparting distinct organoleptic characteristics to the foods, the process of fermentation improves the preservation and safety of easily perishable and highly nutritive raw vegetables through microbial production of organic acids, bacteriocins, alcohols, and other fermentation end products along with transformation of the nutrients in the ingredients (Marco et al., 2021; Ross et al., 2002). Recently, human (Song et al., 2023; Nielsen et al., 2018; Raak et al., 2014; Garnas, 2023) and model animal (Cha et al., 2024; Shahbazi et al., 2021) studies have shown that fermented vegetables can benefit human health. However, the mechanistic details underpinning these outcomes remain to be determined.

Fermented cabbage is a popular fermented vegetable widely consumed across the world and is known by different names (e.g., sauerkraut, suan cai, curtido) (Di Cagno et al., 2016). The basic recipe is essentially mixing shredded cabbage with 2-3% (w/w) sodium chloride and incubating the mixture with minimal exposure to ambient oxygen levels at room temperature for two to three weeks (Medina-Pradas et al., 2017). During a cabbage fermentation, the lack of oxygen and presence of salt selects for lactic acid bacteria (LAB). LAB encompass a group of phylogenetically related bacteria in the Bacillota (formerly Firmicutes) phylum that were traditionally grouped together based on their saccharolytic fermentation energy conservation metabolism resulting in lactic acid as the only or the main (> 50%) end-product (Wang et al., 2021). Culture-dependent and -independent analyses have shown that the dominant LAB in cabbage fermentations change over time, with initial enrichments of heterofermentative species, such as *Leuconostoc mesenteroides* and *Weissella* spp., followed by *Lactiplantibacillus plantarum* and *Levilactobacillus brevis* starting a couple days later (Xu et al., 2024; Gänzle, 2015). It is well-understood that LAB breakdown sugars and other compounds and produce lactic and acetic acid, mannitol, CO_2_, as the primary secreted metabolic end-products, sometimes reaching 2% (v/w) or more in the ferments (Peñas et al., 2017).

Besides these changes, LAB-guided cabbage fermentations have higher concentrations of phenolic compounds (polyphenols, phenolic acids) (Hunaefi et al., 2013; Kaulmann et al., 2014), carotenoids (Kaulmann et al., 2014), glucosinolate breakdown products (ascorbigen, indole-3- carbinol, isothiocyanates) (Krajka-Kuźniak et al., 2011; Martinez-Villaluenga et al., 2012), and other bioactive metabolites (Siddeeg et al., 2022) compared to raw cabbage. Fermented cabbage is also enriched in compounds like phenyl-lactic acid (PLA) (Peters et al., 2019), indole-3-lactic acid (ILA) (Kasperek et al., 2024), and γ-aminobutyric acid (GABA) (Yang, Hu, Xiu, Jiang, Yang, Saren, et al., 2020). These (PLA, ILA, GABA) compounds together with the constituents in fermented cabbage are likely important for its observed immunomodulatory (Huang et al., 2020; Siddeeg et al., 2022; Zubaidah et al., 2020), antioxidant (Krajka-Kuźniak et al., 2011; Xu et al., 2022), and anti-carcinogenic (Peñas et al., 2017; Raak et al., 2014) properties. Although only one human study on fermented cabbage has been performed, it was found that both pasteurized and unpasteurized sauerkraut significantly improved irritable bowel syndrome (IBS) symptoms (*p* < 0.04) (Nielsen et al., 2018).

Of particular interest is the capacity of fermented cabbage and related foods to alter intestinal barrier function. The intestinal epithelium serves as a selectively permeable barrier that allows the absorption of fluids and nutrients while limiting the translocation of microorganisms, antigens, and other toxic compounds into the sterile serosal circulation of the host from the gut lumen (Odenwald & Turner, 2017; Lee et al., 2018). Compounds, such as cytokines, growth hormones, dietary components, microbial molecules, influence intestinal barrier integrity and function through modulating junctional complexes (Odenwald & Turner, 2017). For instance, the proinflammatory cytokine tumor necrosis factor alpha (TNF-α) increases the expression of myosin light chain kinase (MLCK), which then causes the re-organization of the peri-junctional actin and tight junction proteins (e.g., occluding, ZO-1), resulting in increased paracellular permeability (Lee et al., 2018). *In vitro* cell line models (e.g., Caco-2, T84, HT-29) (Schoultz & Keita, 2020) have played a key role in facilitating the functional and mechanistic understanding of the association between disrupted intestinal barrier integrity and numerous inflammatory and metabolic diseases (e.g., inflammatory bowel disease, obesity) and therapeutic development (Odenwald & Turner, 2017; Stolfi et al., 2022).

To better understand the potential of fermented cabbage to impact intestinal function, we evaluated cell-free preparations of cabbage and commercial and laboratory-scale fermented cabbage (sauerkraut) ferments. Homogenates of fermented cabbage prepared from an estimated serving size of 10 gram were tested for their capacity to protect against pro-inflammatory cytokine (IFN-γ and TNF-α)-induced damage in a polarized Caco-2 cell model. Some laboratory fermentations also incorporated the use of a probiotic strain of *L. plantarum* NCIMB8826R (LP8826R). This strain was shown to increase localization of the tight junction protein ZO-1 in healthy individuals (Karczewski et al., 2010) as well as induce intestinal epithelial repair (Crakes et al., 2019; Hirao et al., 2014) and exert anti-inflammatory (Yin et al., 2018) and anti-obesogenic (Heeney et al., 2018) effects in animal models. To begin to understand the differences between cabbage and fermented cabbage, metabolites were quantified with (un)targeted metabolomics using gas chromatography-time of flight mass spectrometry (GC-TOF/MS) and reverse-phase liquid chromatography high resolution tandem mass spectrometry (RP-LC-HRMS/MS). Based on those findings, lactate, D-PLA, and ILA, were investigated for barrier-protective capacities in the polarized Caco-2 cell model.

## Materials and Methods

### Bacterial strain growth conditions and preparation of inoculum for cabbage fermentations

Isolation of the rifampicin-resistant mutant *L. plantarum* NCIMB8826R (LP8826R) was described previously (Yin et al., 2018). LP8826R was routinely grown at 37 °C in a commercial preparation of lactobacilli de Man, Rogosa and Sharpe (MRS) medium (Becton-Dickinson, Franklin Lakes, NJ, USA). When appropriate, 50 µg/ml rifampicin (Thermo Fisher Scientific, Waltham, MA, USA) was added to the MRS for LP8826R enumeration. For inoculation into sauerkraut fermentations, LP8826R was incubated in MRS for 24 h, collected by centrifugation at 9,000 x g for 3 min at room temperature, washed twice in sterile phosphate buffered saline (PBS; 137 mM NaCl, 2.7 mM KCl, 4.3 mM Na_2_HPO_4_*7H_2_O, 1.4 mM KH_2_PO_4_, pH 7.2), and then suspended in sterile PBS.

### Laboratory-scale fermentations (LSF)

Green cabbages (*Brassica oleracea* var. capitata), purchased from local grocery stores, were shredded using a grater, mixed with 2% canning salt (w/w) (Morton Salt, Chicago, IL, USA), dispensed into 28 oz sterile WhirlPak bags with a 330 µm filter (Whirl-Pak, Pleasant Prairie, WI, USA), and covered with a 2% (w/w) brine solution of canning salt until all pieces were submerged. The mixtures were divided into six containers and incubated at 22 °C. After 3 days, LP8826R in 1.75 to 2 ml PBS was added to half of the containers (n = 3) to reach 5 x 10^7^ CFU/g. The same volume of PBS was added to the other laboratory ferments. The LSFs were performed twice, once with a total fermentation time of 14 days and the other for 21 days. Brine pH (SevenEasy pH meter (Mettler-Toledo, Columbus, OH, USA) and salinity (SPER Scientific, Scottsdale, AZ, USA) were measured for the first seven days and then at the end of the study.

### Commercial sauerkraut

Containers (n = 6) of commercial sauerkraut from the same brand were purchased from local supermarkets between December 2021 and October 2022 (**Table 1**). They were stored at 4 °C and sampled within 3 days of purchase.

**Table 1.** Parameters of commercial sauerkraut sampled.

| **LOT #** | **Best By Date** | **Sampling Date** | **Brine Salinity (ppt)** | **Homogenate pH** | **LAB (log<sub>10</sub> CFU/ml)<sup>a</sup>** |
| --- | --- | --- | --- | --- | --- |
| ND | ND | Dec 08, 2021 | ND | 3.37 | ND |
| ND | ND | Nov 12, 2021 | ND | 3.20 | ND |
| 2022234 | 012623 | Sep 28, 2022 | 26.7 | 3.37 | 6.60 |
| 2022209 | 122922 | Sep 28, 2022 | 27.5 | 3.30 | 6.95 |
| 2022234 | 012623 | Sep 28, 2022 | 27.4 | 3.36 | 6.99 |
| 2022166 | 111922 | Oct 09, 2022 | 25.3 | 3.42 | 1.90 |

ND: not determined.
<sup>a</sup> Estimated on MRS laboratory culture medium

### Intestinal epithelial cell culture maintenance

Caco-2 cells ATCC HTB-37 were purchased from the American Type Cell Culture (Rockville, MD, USA) and grown in DMEM as previously described (Hubatsch et al., 2007). The cell culture reagents, Dulbecco’s Modified Eagle Medium (DMEM), fetal bovine serum (FBS), sodium pyruvate (100X), nonessential amino acids (100X), GlutaMax (100X), penicillin/streptomycin 10,000 U/ml, Trypsin-EDTA (0.25%, 1X), and sterile Dulbecco’s phosphate buffer saline (D-PBS, pH 7.4, no calcium, no magnesium), were obtained from Gibco (Life Technologies, Carlsbad, CA, USA). Briefly, the Caco-2 cells were incubated in DMEM containing 20% FBS, 1 mM sodium pyruvate, 0.1 mM nonessential amino acids, 2 mM GlutaMax, and 1% v/v penicillin/streptomycin at 37 °C in 10% CO_2_, split using Trypsin-EDTA, and passaged in 75 cm^2^ culture flasks (Fisher Scientific, Waltham, MA, USA) until they were ready for seeding.

### Preparation of (fermented) cabbage homogenates for bacterial enumeration and application onto Caco-2 monolayers

Approximately 10 g cabbage (either fermented or raw) and 1 ml brine were collected from the same container and homogenized in sterile WhirlPak bags with 330 µm filter using a rubber mallet. The homogenates were then serially diluted in PBS and plated on MRS agar containing 25 µg/ml natamycin (Dairy Connection, WI, USA), an antifungal compound, for total bacterial cell counts and MRS agar containing both 25 µg/ml natamycin and 50 µg/ml rifampicin for LP8826R enumeration.

For experiments with Caco-2 cells, cabbage homogenates were centrifuged at 10,000 x g for 10 min at 4 °C. The supernatant was collected and adjusted to a pH of 7.4 using either 1M or 6M sodium hydroxide (NaOH) prior to passing through a 0.22 µm surfactant-free cellulose acetate (SFCA) filter (Globe Scientific, Mahwah, NJ, USA). The collected cell-free supernatants from LSF after 7 days of fermentation (n = 3) and commercial cabbage ferment (n = 1) were then diluted to 1%, 10%, or 50% v/v with Caco-2 cell culture medium (DMEM) containing 1 mM sodium pyruvate, 0.1 mM nonessential amino acids, and 2 mM GlutaMAX to determine the appropriate concentration for application to Caco-2 monolayers. The concentration of cell-free preparation of cabbage homogenates in Caco-2 cell culture medium at 10% v/v were selected after this initial test. A 2.91% (w/v) solution of NaCl was made to match the salinity of commercial fermented cabbage homogenate after pH adjustment (29.1 ppt) and used to examine the impact of salt on the Caco-2 cell monolayers. The salt solution was then diluted to 10% (v/v) in DMEM containing supplements (1 mM sodium pyruvate, 0.1 mM nonessential amino acids, 2 mM GlutaMax). The prepared homogenates and salt solution were stored at -80 °C until use.

### Lactate, ILA, and D-PLA preparation for application onto Caco-2 monolayers

DL-lactic acid (W261106-1KG-K), DL-indole-3-lactic acid (ILA; I5508-250MG-A), D- (+)-3-Phenyllactic acid (D-PLA; 376906-5G) were obtained from Sigma-Aldrich (St. Louis, MO, USA). Lactate and D-PLA were resuspended in sterile water and ILA was mixed with 0.052% v/v DMSO. All three compounds were prepared in concentrations that were 10-fold higher than intended for use in cell culture. To ensure D-PLA and ILA were dissolved, the solutions were sonicated for 10 min at 40 kHz in a Branson 8510 Ultrasonic Cleaner bath (Branson Ultrasonics Corp., Danbury, CT, USA). The metabolite solutions were then adjusted to a pH of 7.40 ± 0.04 with NaOH and diluted in DMEM containing 1 mM sodium pyruvate, 0.1 mM nonessential amino acids, and 2 mM GlutaMax to reach a final concentration of 50 mM lactate, 60 µM D- PLA, and 25 µM ILA (0.0052% (v/v) DMSO), both separately and combined. The DMEM suspensions were sterilized using a 0.22 µm SFCA filter and stored at -80 °C until use.

### Intestinal epithelial monolayer cytokine stimulation

To form a monolayer, Caco-2 cells between passages 35 and 36 were seeded onto polycarbonate membranes in Transwell inserts (0.33 cm^2^ surface area, 6.5 mm, 0.4 µm pore size, 24 wells; Corning, NY, USA) at a density of 1 x 10^4^ cells/cm^2^. After incubation for at least 21 days in DMEM containing 1 mM sodium pyruvate, 0.1 mM nonessential amino acids, and 2 mM GlutaMax, the cells were fully differentiated and formed polarized monolayers with reduced permeability as quantified by trans-epithelial electrical resistance (TER). TER was measured after 15 min equilibration at room temperature with an epithelial voltohmmeter (World Precision Instruments, Sarasota, FL, USA) equipped with a STX-2 “chopstick” electrode. Caco-2 monolayers with an initial TER of more than 250 Ohm·cm^2^ were used for the subsequent experiments.

Prior to the application of either the (fermented) cabbage homogenates or compounds, TER values were measured. DMEM supplemented with 1 mM sodium pyruvate, 0.1 mM nonessential amino acids, 2 mM GlutaMax and containing either cabbage homogenates (10% v/v), selected metabolites (D-PLA, ILA, lactate, or the three combined), sterile water (10% v/v) or 0.0052% DMSO were applied to the apical side of the monolayers. After 3 h incubation at 37 °C, 100 ng/ml IFN-γ (R&D Systems, Minneapolis, MN, USA) was added to the basolateral compartment. The medium in the basolateral compartment was replaced with DMEM with 1 mM sodium pyruvate, 0.1 mM nonessential amino acids, 2 mM GlutaMax supplemented with 10 ng/ml TNF-α (R&D Systems, Minneapolis, MN, USA) 21 h later. TER values of the monolayers were measured immediately prior to TNF-α exposure and 24, 32, and 48 h later.

### Paracellular permeability assay

Paracellular permeability was determined by the flux of fluorescein isothiocyanate-dextran (FITC; 4 kDa; Sigma-Aldrich, St. Louis, MO, USA) from the apical to basolateral side of differentiated Caco-2 monolayers as previously described (Zhai et al., 2019). After 48 h stimulation with TNF-α, apical DMEM was replaced with FITC (2 mg/ml) in DMEM with 1 mM sodium pyruvate, 0.1 mM nonessential amino acids, 2 mM GlutaMax. After 30, 60, 90, and 120 min, FITC concentrations on the basolateral side were quantified by measuring the fluorescence intensity (485 nm excitation and 530 nm emission) in a microplate reader (Synergy-2, BioTek, Winooski, VT, USA). FITC levels were calculated using a standard curve of FITC in DMEM with the indicated supplements. The apparent permeability coefficient, P_app_ (cm/s), was determined using the following equation: P_app_ = [dQ/dt] x [1 / (A x C_0_)], where dQ/dt is the quantity of FITC transported per second (ng/sec), A is the surface area of the filter (cm^2^), and C_0_ is the initial FITC concentration in DMEM on the apical side (ng/ml).

### ELISA quantification of IL-8

After 48 h stimulation with TNF-α, spent media collected from the basolateral compartment was used for the quantification of interleukin 8 (IL-8) via ELISA (Life Technologies, Carlsbad, CA, USA) according to the manufacturer’s instructions.

### RNA extraction and reverse-transcription quantitative PCR (RT-qPCR)

After 48 h exposure to TNF-α, total RNA from the Caco-2 monolayers was extracted using TRIzol (Invitrogen, Carlsbad, CA, USA) and purified using the TRIzol-chloroform protocol. RNA was quantified on the NanoDrop 2000c (Thermo Fisher, Waltham, MA, USA) and found to be intact (RIN 8.8 - 9.8) according to the Agilent 2100 Bioanalyzer (Agilent Technologies, Santa Clara, CA, USA). RNA was reverse transcribed to cDNA using the High-Capacity cDNA Reverse Transcription kit (Thermo Fisher Scientific, Waltham, MA, USA). RT-qPCR was performed using the Fast SYBR Green Master Mix (Thermo Fisher Scientific, Waltham, MA, USA) and 200 nM of the designated primers (**Table 2)** on a 7500 Fast Real-time PCR system (Applied Biosystems, Carlsbad, CA, USA). Amplification was initiated at 95 °C for 20 sec and was followed by 40 cycles of 95 °C for 3 sec and 60 °C for 30 sec. Primer specificity was assessed using melting curves following the amplification stage. All reactions were performed in duplicate and reactions lacking RT controls for each sample were included to check for DNA contamination. Data were normalized using the 2^-ΔΔCt^ method with *GAPDH* as the housekeeping gene and untreated cells as the reference condition.

**Table 2.**
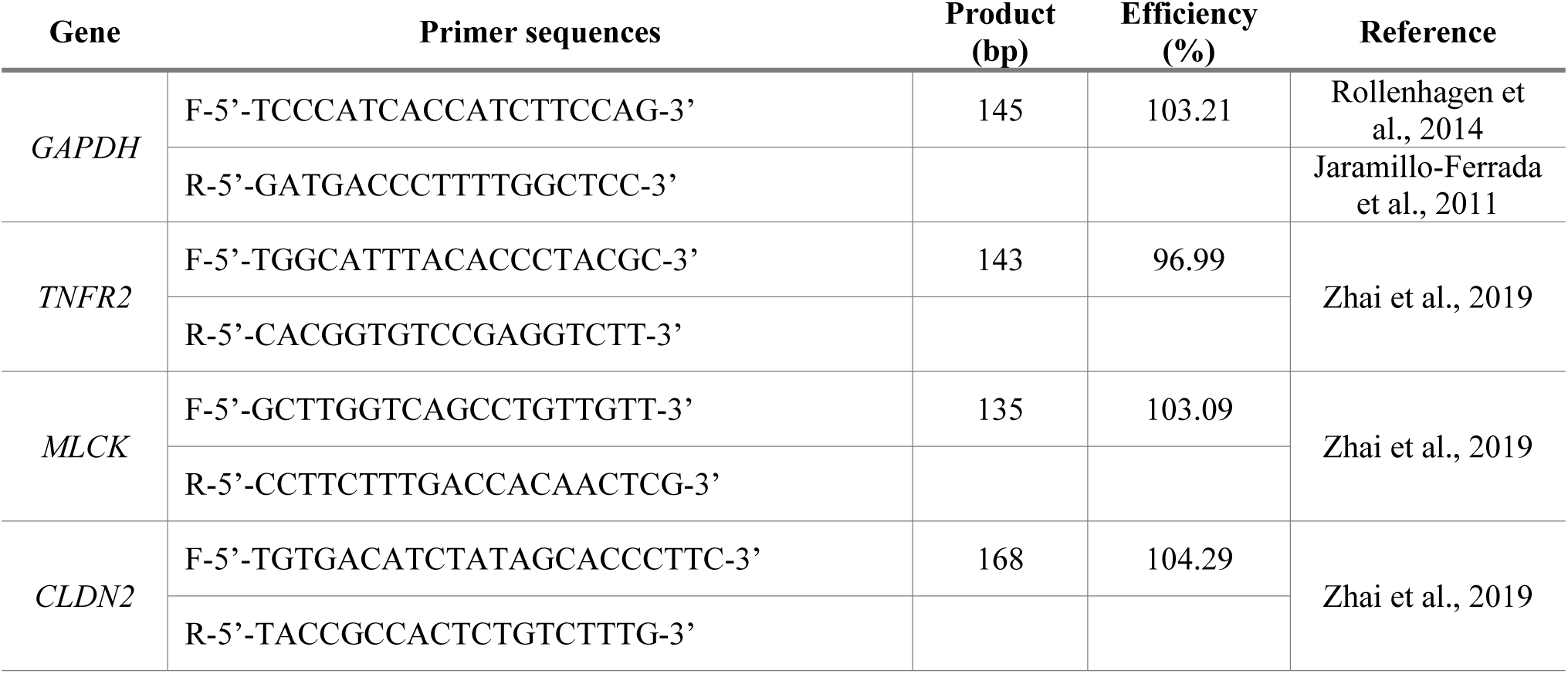
qRT-PCR primer DNA sequences.

### Metabolomic analysis of cell-free (fermented) cabbage homogenates

Homogenates from the LSF and commercial ferments were prepared for GC-TOF/MS and RP-LC-HRMS/MS at the UC Davis West Coast Metabolomics Center (WCMC) (https://metabolomics.ucdavis.edu/) as described previously (Fiehn, 2016; Folz et al., 2023). A complete list of detectable metabolites using these methodologies can be found at https://metabolomics.ucdavis.edu/metabolites (**Supplemental File 1**).

GC-TOF/MS analysis was performed on an Agilent 7890A gas chromatograph (Agilent, Böblingen, Germany) coupled to a Leco Pegasus IV TOF mass spectrometer (Leco, St. Joseph, MO, USA). Raw untargeted GC-TOF/MS data files were processed by Leco ChromaTOF software and further analyzed using the BinBase pipeline (Skogerson et al., 2011) for identifying metabolites by retention index matching and mass spectra alignment. The method was designed for untargeted and targeted primary metabolite detection and quantification, namely for carbohydrates and sugar phosphates, amino acids, hydroxyl acids, free fatty acids, purines, pyrimidines, and aromatics (Fiehn, 2016). The levels of ILA, PLA, and GABA in (fermented) cabbage homogenates were quantified through targeted GC-TOF/MS.

RP-LC-HRMS/MS was performed using a Vanquish LC (Thermo Scientific, Waltham, MA, USA) coupled to a Q-Exactive HF^+^ orbital ion trap mass spectrometer (Thermo Scientific) with positive and negative ion electrospray ionization (ESI) modes. MS-DIAL v. 4.92 (Tsugawa et al., 2020) was used to process untargeted LC-MS/MS data with retention times and MS/MS spectra matching compared to the MassBank of North America (Horai et al., 2010). Using this method, flavonoids, anthocyanins, polyphenols, alkaloids, coumarins, terpene-and polyphenol-glycosides, kaempferols, flavanones, cinnamates, and related compounds were detected.

Information on data acquisition, processing, and raw data normalization can be found in the document shared by the WCMC (**Supplemental File 2**). As described in **Supplemental File 2**, untargeted raw metabolite chromatogram data were normalized to the identified metabolite total ion chromatogram (mTIC), which is the sum of all peak heights for all identified metabolites (excluding the unknowns) in each sample. For targeted GC-TOF/MS, PLA, ILA, and GABA were normalized to the amount of sample injected onto the column. Untargeted GC-TOF/MS data were provided by WCMC as a table containing identified BinBase name, peak heights, retention index, m/z ratio, PubChem ID, KEGG ID, and InChl Key values. RP-LC-HRMS/MS data were provided by WCMC as a table containing identified metabolite name, MS-DIAL identifier, peak heights, m/z ratio, ESI mode, InChI key and retention time.

For statistical analysis, peak height datasets were uploaded to MetaboAnalyst 6.0 (Pang et al., 2024; Xia & Wishart, 2011). Features (metabolites) with > 50% missing values were removed, and the remaining missing values were replaced using the Limit of Detection (LoD) method (one fifth of the minimum positive value of each variable) (Sun & Xia, 2024).

Furthermore, features with > 10% inter-quantile range (IQR) variance were filtered out. As noted previously (Folz et al., 2023), for GC-TOF/MS data, the peak heights were log_10_ transformed prior to analysis. For the RP-LC-HRMS/MS data (Folz et al., 2023), peak heights were normalized to the sums of the peak heights of internal standards (1-cyclohexyl-dodecanoic acid urea (CUDA), D3-L-Carnitine, D4-Daidzein, D5-Hippuric acid-d5, D9-Reserpine, Val-Tyr-Val) (**Supplemental File 3**) added to each sample and then subjected to log_10_ transformation.

### Statistical analysis

Data were analyzed using GraphPad Prism 10 software for Windows (GraphPad Software, Inc., La Jolla, CA, USA), unless stated otherwise. The cabbage fermentation (pH, bacterial cell counts), Caco-2 cell culture (TER, FITC, IL-8, RT-qPCR), and targeted GC-MS data were reported as mean ± standard deviation (SD) and analyzed by ANOVA with post hoc test for comparing multiple treatments.

For untargeted metabolomics datasets, principal component analysis (PCA) was applied to evaluate the compositions of the six different (fermented) cabbage homogenates. To identify metabolites significantly elevated or reduced after fermentation, the relative peak heights of fermented homogenates were compared to the raw cabbage homogenates (D0) using the Wilcoxon rank-sum test (FDR of 0.05) (**Supplemental Files 4** and **5**). Heatmaps of the top 35 significantly changed metabolites with the lowest *p* values were generated via hierarchical cluster analysis (Euclidean distance measure, Ward clustering method). PERMANOVA and pairwise Adonis tests were performed in the LMSstat package (https://github.com/CHKim5/LMSstat/blob/master/README.md) and the pairwiseAdonis R package (Martinez Arbizu, 2020), respectively, in R 4.3.0 (R Core Team, 2023).

## Results

### LP8826R dominates and modifies cabbage fermentations

According to culturable bacterial cell numbers on MRS agar, a nutritionally replete medium used to enrich LAB (De Man et al., 1960), there were 3.68 ± 3.55 log_10_ CFU/ml in the cabbage homogenate prior to the start of the laboratory scale fermentations (LSF). This number increased to 9.19 ± 8.91 log_10_ CFU/ml after two days of incubation at 22 °C (**Fig 1A**). Bacterial numbers then declined by approximately 10-fold to approximately 8.0 log_10_ CFU/ml by day 4 and were sustained at that level for at least 10 days. By day 21 of the LSF, average cell numbers declined by another 2-fold to 7.61 ± 7.82 log_10_ CFU/ml homogenate (**Fig. 1A**).

**Figure 1.**
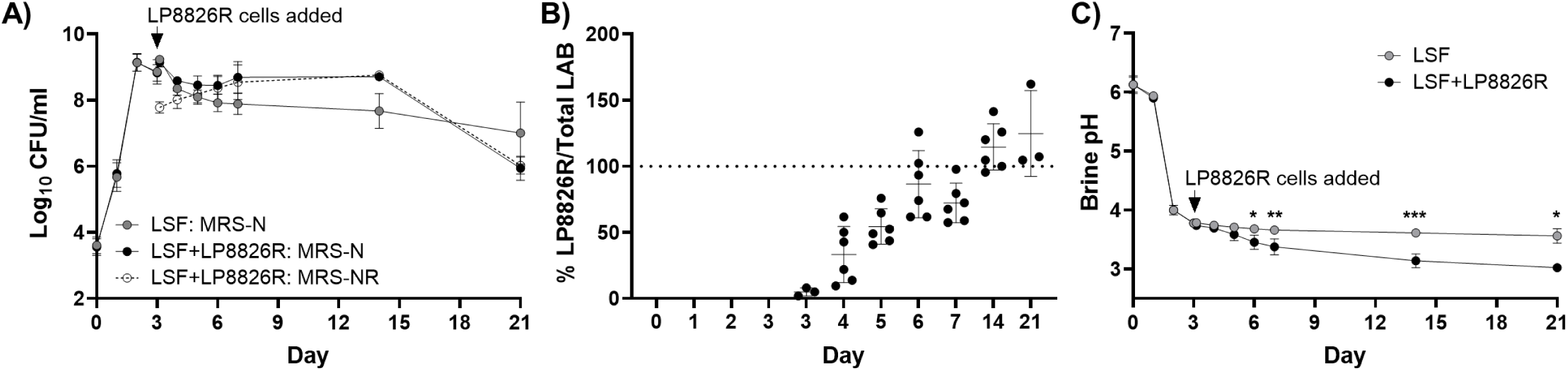
Bacterial numbers and brine pH in laboratory-scale ferments (LSF). (**A**) LAB in LSF (gray circles), LAB in LSF with LP8826R (black circles), and LP8826R in the LSF with that added strain (open circles). LAB were enumerated on MRS with natamycin (25 µg/ml) (MRS-N) and LP8826R was enumerated on MRS containing natamycin (25 µg/ml) and rifampicin (50 µg/ml) (MRS-NR). Rifampicin resistant bacteria were not detected in the LSF lacking LP8826R additions (below 3.33 x 10^1^ CFU/ml limit). (**B**) The percentage of LP8826R to total LAB numbers over time, and (**C**) brine pH. The mean ± SD is shown for 6 replicates. To determine differences in brine pH, two-way mixed effect ANOVA with Tukey’s multiple comparisons test was performed (* p < 0.05; ** p < 0.01; *** p < 0.001).

Addition of LP8826R to the LSF on day 3 briefly elevated the total culturable LAB amounts in cabbage fermentations from 8.88 ± 8.62 log_10_ CFU/ml homogenate before inoculation to 9.15 ± 8.65 log_10_ CFU/ml approximately 3 h after inoculation (**Fig. 1A**). Over the subsequent three days, the numbers of LP8826R increased approximately 5-fold to 8.49 ± 8.35 log_10_ CFU/ml, as estimated on MRS with rifampicin (**Fig. 1A**). On that day, LP8826R constituted 86.50 ± 25.43% of total culturable LAB in the ferments (**Fig. 1A** and **Fig. 1B**). This percentage was sustained until the end of the study (**Fig. 1B**). Compared to the LSF, the total culturable LAB numbers in the LSF + LP8826R ferments were significantly higher on day 4 (*p* = 0.0320), day 5 (*p* = 0.0374), day 6 (*p* = 0.0298), and day 14 (*p* < 0.0001) (**Fig. 1A**). At day 21, total culturable LAB numbers between the LSF and LSF + LP8826R were not significantly different (*p* = 0.4015) (**Fig. 1A**).

In both LSF and LSF + LP8826R, reductions in pH coincided with the increase in bacterial numbers. The pH declined from pH 6.13 ± 0.13 at the start of the study to pH 4.00 ± 0.08 after the first 2 days of incubation (**Fig. 1C**). For the LSF, the brine pH continued to decrease, reaching pH of 3.61 ± 0.03 and 3.56 ± 0.12 on day 14 and 21, respectively (**Fig. 1C**). Brine pH values were significantly lower in the LSF + LP8826R compared to the LSF starting on day 6 of the fermentation (*p* < 0.05) (**Fig. 1C**). The significant reduction (*p* = 0.0206) in culturable LAB numbers between day 14 and day 21 observed in LSF + LP8826R may have been the result of the lower pH in those fermentations.

Brine salinity was measured at days 14 and 21 of the study. Brine salinity in the LSF was similar at both time points (19.90 ± 1.57 parts per thousand (ppt)) on day 14) and 19.50 ± 0.17 ppt on day 21). The addition of L8826R did not result in changes to salinity on day 14 (20.53 ± 1.76 ppt) and 21 (20.17 ± 0.45 ppt) compared to LSF without this strain.

Commercial sauerkraut contained 6.54 ± 6.66 log_10_ CFU/ml homogenate according to growth on MRS agar (**Table 1**). These bacterial numbers were similar to levels found in the LSF on days 14 and 21 and in LSF + LP8826R on day 21, but less than those detected in LSF + LP8826R on day 14 (*p* < 0.0001). The pH of commercial sauerkraut homogenate (pH 3.34 ± 0.08) was significantly lower compared to LSF on days 14 and 21, but significantly higher compared to the LSF + LP8826R ferments (*p* ≤ 0.0092) (**Fig. 1C** and **Table 1**). The salinity of commercial sauerkraut brine was 26.73 ± 1.01 ppt, a value significantly higher compared to the LSF and LSF + LP8826R (*p* < 0.002) (**Table 1**).

### Cabbage ferments prevent cytokine-induced intestinal barrier disruption

To establish a suitable range of fermented cabbage homogenate for assessment on Caco-2 cell monolayers, Caco-2 cell culture medium (DMEM) was adjusted to contain 1, 10, or 50% (v/v) of filtered, pH-adjusted (pH 7.4) homogenates from day 7 LSFs and the commercial cabbage ferment. Compared to untreated controls, the apical application of either 1 or 10% (v/v) of the homogenates did not significantly change TER or permeability to FITC-dextran (FITC) with sequential basolateral exposure to IFN-γ for 21 h and TNF-α for another 48 h (data not shown). However, TER was negatively affected when 50% (v/v) was applied, resulting in an average 10-fold reduction (data not shown). This change was not a result of diluting the DMEM culture medium since applying an equal quantity of water to the controls did not result in TER reductions or increases in paracellular permeability to FITC (data not shown). Because there were no significant differences in TER and FITC permeability for the Caco-2 cell monolayers treated with either 1 or 10% v/v of the fermented cabbage homogenate in DMEM, 10% (v/v) of the pH-adjusted and filtered cabbage and sauerkraut homogenates were investigated further.

As found previously (Zhai et al., 2019), basolateral incubation with IFN-γ for 21 h followed by TNF-α resulted in significant reductions of TER compared to untreated controls within 24 h after application that were sustained for the duration of the experiment (**Fig. 2A** and **Fig. S1;** *p* < 0.0001). IFN-γ and TNF-α induced reductions in TER were not found for monolayers incubated with the LSF, irrespective of time of sampling or LP8826R, and instead TER levels were similar to the untreated controls (*p ≥* 0.2008) (**Fig. 2A**). TER of Caco-2 monolayers exposed to the fermented cabbage homogenates were also significantly greater than the IFN-γ and TNF-α treated controls, with exception of the LSF collected at day 14 and measured 24 h after TNF-α addition (**Fig. S1**). By comparison, raw cabbage homogenates (cabbage in brine from day 0 of the study) had reduced TER compared to the controls at all time points (**Fig 2A** and **Fig S1**) and were not significantly different from the cytokine-treated controls starting 32 h after TNF-α application (**Fig. S1**). To investigate effects of cabbage and saline on TER, Caco-2 cell monolayers were either treated with cabbage homogenate collected after homogenization in water (instead of saline) or DMEM adjusted to contain 10% (v/v) of a 29.1 ppt NaCl brine solution prepared to match the salinity of the fermented cabbage. The former preparation did not result in barrier protection, as indicated by the significantly lower TER (*p* ≤ 0.03) compared to untreated controls (**Fig. S2A**). The addition of NaCl to DMEM resulted in no significant differences in TER compared to either the IFN-γ and TNF-α treated or untreated controls (**Fig. S2A**).

**Figure 2.**
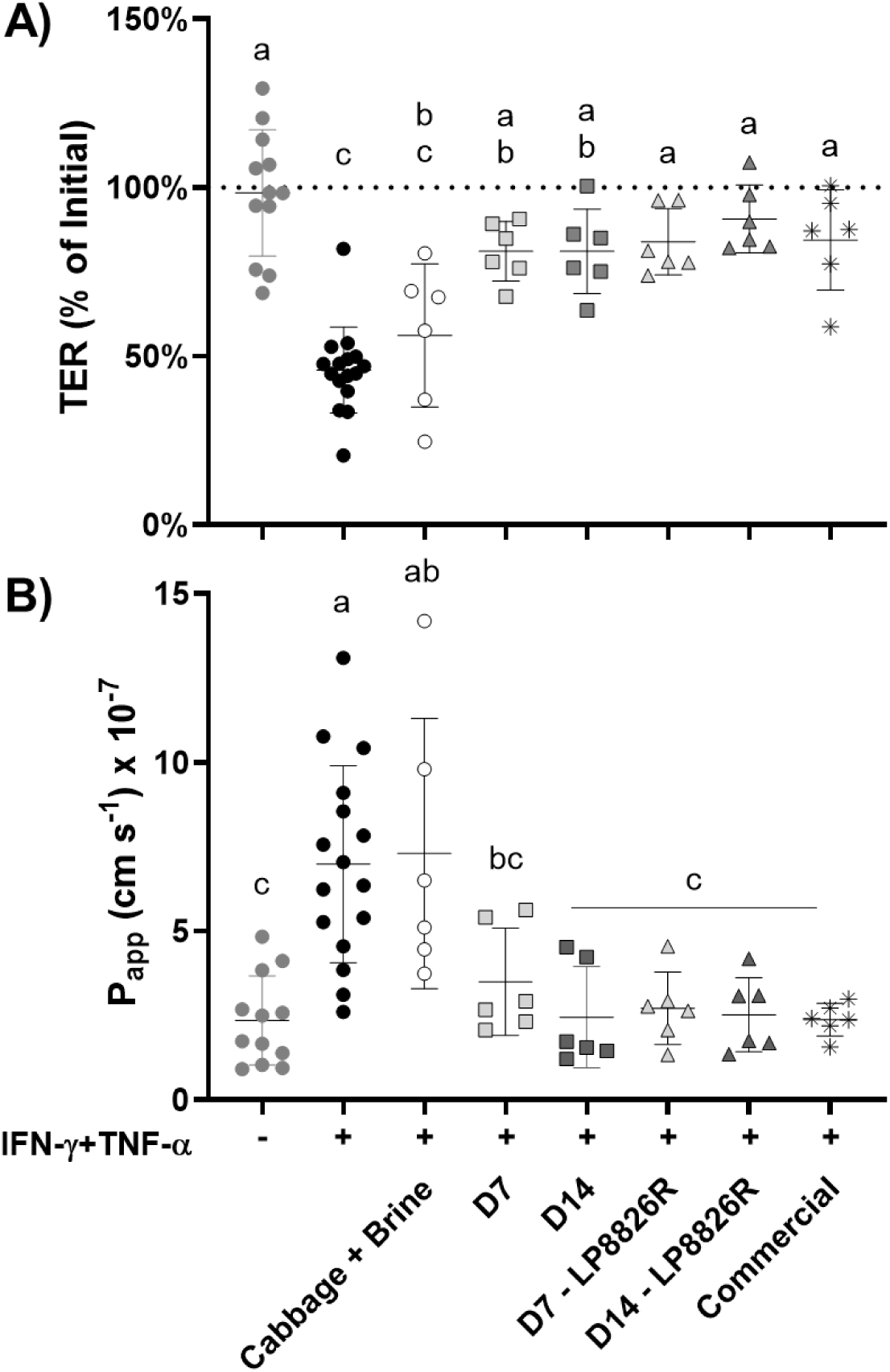
Effect of (fermented) cabbage homogenates on intestinal barrier permeability of cytokine-perturbed Caco-2 monolayers. (**A**) Trans-epithelial electrical resistance (TER) of Caco-2 monolayers 48 h after basolateral IFN-γ and TNF-α sequential additions. Values are normalized to TER immediately prior to when TNF-α was added. (**B**) Apparent permeability coefficient (P_app_) to FITC-dextran (4kD) of Caco-2 monolayers at the 48 h time point. D7 and D14 were collected on days 7 and 14, respectively, from LSF either with or without the LP8826R additions. Replicates included Caco-2 monolayers not exposed to IFN-γ and TNF-α (n = 12), controls to which the cytokines were applied (n = 16), and those exposed to the cytokines and (fermented) cabbage (n = 6). The mean ± SD is shown for four independent experiments. The letters indicate significant differences based on one-way ANOVA with Tukey’s multiple comparisons test.

Assessments of paracellular permeability to FITC-dextran were performed 48 h after TNF-α addition. Significantly higher quantities of FITC were detected in the basolateral compartment of Caco-2 cell monolayer controls treated with IFN-γ and TNF-α compared to those not exposed to the cytokines (**Fig. 2B**; P_app_ coefficient; *p* < 0.0001), confirming that the cytokines disrupted barrier integrity. Notably, Caco-2 monolayers to which either LSF, LSF + LP8826R, or the commercial product homogenate were added had significantly lower permeability to FITC (*p* < 0.0292) than the cytokine-treated controls and were equal to the untreated controls (**Fig. 2B**). In contrast, FITC translocation levels increased for Caco-2 monolayers exposed to the raw cabbage homogenates (D0) (*p* < 0.0007) (**Fig. 2B**). Caco-2 cell monolayers treated with cabbage homogenate collected after homogenization in water (instead of saline) also had an elevated permeability to FITC (*p* = 0.0003) compared to untreated monolayers (**Fig. S2B**). Lastly, Caco-2 cell monolayers exposed to NaCl (2.91 ppt) had intermediate levels of FITC translocation that were not significantly different from either the untreated and cytokine, IFN-γ and TNF-α, treated monolayers (**Fig. S2B**). Together, these results indicate that fermented cabbage, but not raw cabbage or brine, had the capacity to protect against inflammation-induced intestinal barrier damage.

### Fermented cabbage variably altered IL-8 secretion and expression of barrier-modulatory genes

Concentrations of IL-8, a pro-inflammatory chemokine and indicator of inflammatory responses in epithelial cells (Jobin et al., 1999), were significantly higher in the basolateral medium of Caco-2 cell monolayers exposed to IFN-γ and TNF-α compared to the untreated controls (*p* < 0.0001) (**Fig. 3**). Similarly high levels of IL-8 were found for the LSF with and without LP8826R (**Fig. 3**). Quantities of IL-8 were significantly reduced for monolayers exposed to the commercial product compared to controls exposed to IFN-γ and TNF-α (*p* = 0.0138), albeit the quantities were still higher than the controls not exposed to the cytokines (**Fig. 3**). Interestingly, the levels of IL-8 were similarly reduced after exposure to the raw cabbage and brine mixture (D0) (**Fig. 3**).

**Figure 3.**
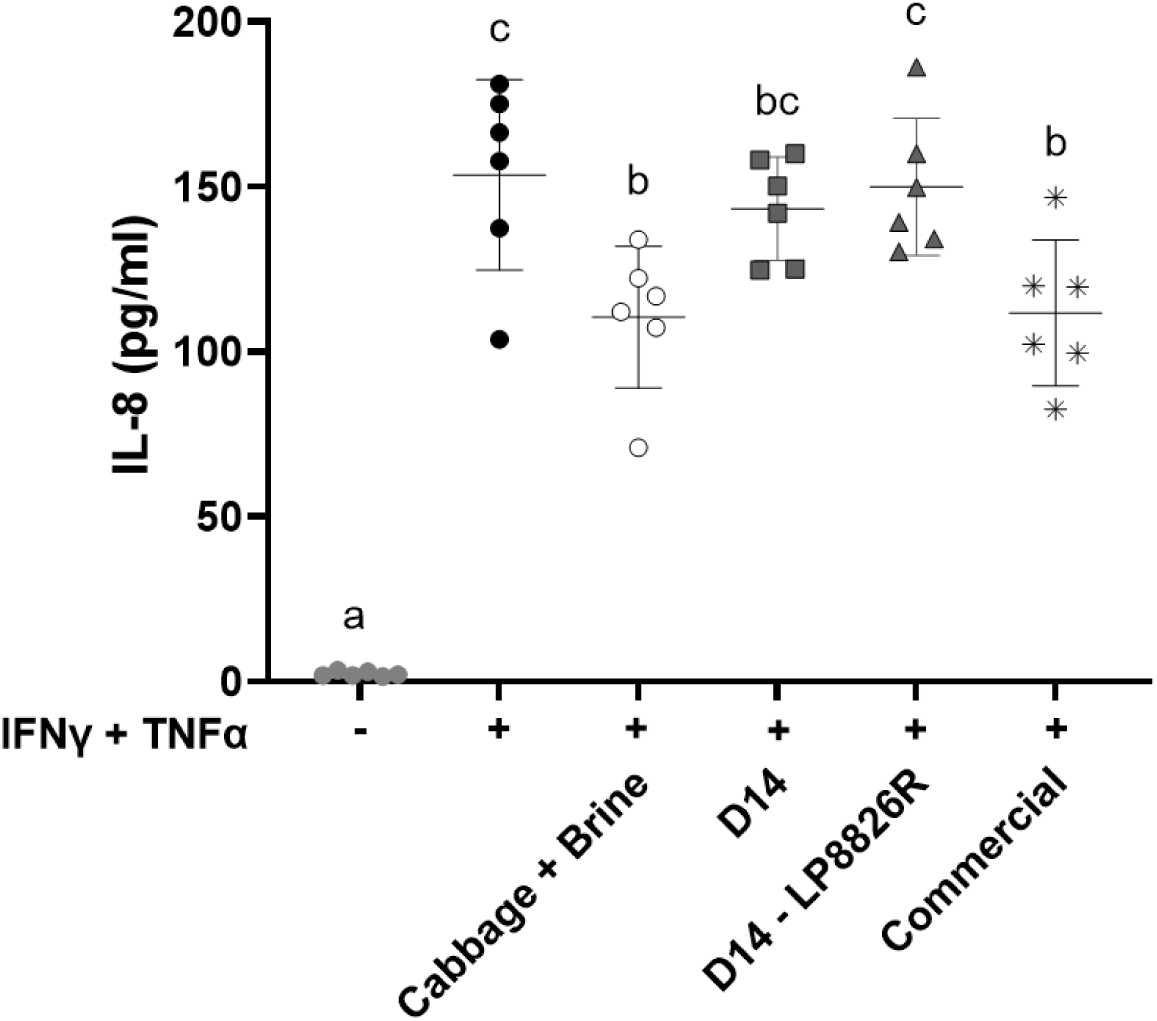
IL-8 production by Caco-2 monolayers after sequential exposure to IFN-γ and TNF-α. DMEM was collected from the basolateral side of Caco-2 monolayers 48 h after sequential application of IFN-γ and TNF-α. D14 and D14 – LP8826R were collected on day 14 from LSF either without or with LP8826R addition. The mean ± SD is shown for selected basolateral spent media from four independent experiments (n = 6). Letters indicate significant differences based on one-way ANOVA with Tukey’s multiple comparisons test.

To investigate the effects of the fermented cabbage on Caco-2 gene expression, RT-qPCR was performed to quantify levels of the TNF receptor (*TNFR2*), myosin light chain kinase (*MLCK*), and *CLDN2*, encoding for the pore-associated tight junction protein Claudin-2. Transcript levels of *TNFR2* and *CLDN2* were approximately 2-fold higher with IFN-γ and TNF- α exposure compared to the controls. Surprisingly, *MLCK* expression was slightly reduced by cytokine challenge (**Fig. S3**). Application of fermented cabbage homogenates prior to IFN-γ and TNF-α did not significantly alter the expression levels of *TNFR2*, *MLCK*, or *CLDN2*, with the exception for the LP8826R-inoculated cabbage ferments, which had similar *MLCK* gene expression as found for the controls (**Fig. S3**).

### Raw and fermented cabbage homogenates have distinct metabolomes

GC-TOF/MS detected 583 metabolites in (fermented) cabbage homogenates, among which 147 were identified (**Supplemental File 4**). Principal component analysis (PCA) showed that the metabolome profiles of the raw and fermented homogenates were significantly different (PERMANOVA *p* = 0.001, R_2_ = 0.95305) (**Fig. 4A**). Metabolites in the raw cabbage (D0) clustered separately from the fermented cabbage homogenates (*p_adj_* ≤ 0.06, pairwise Adonis test) (**Fig. 4A**). The commercially prepared ferments were also distinct from the LSFs (*p_adj_*= 0.045, pairwise Adonis test) (**Fig. 4A**). Among the LSFs, those with LP8826R were similar to the LSF prepared without this exogenous strain (*p_adj_* = 0.210) on day 7 of the study, but then changed and were significantly different from the LSFs on day 14 (*p_adj_*= 0.045) and resembled the commercial sauerkraut (*p_adj_* = 0.075, pairwise Adonis test) (**Fig. 4A**).

**Figure 4.**
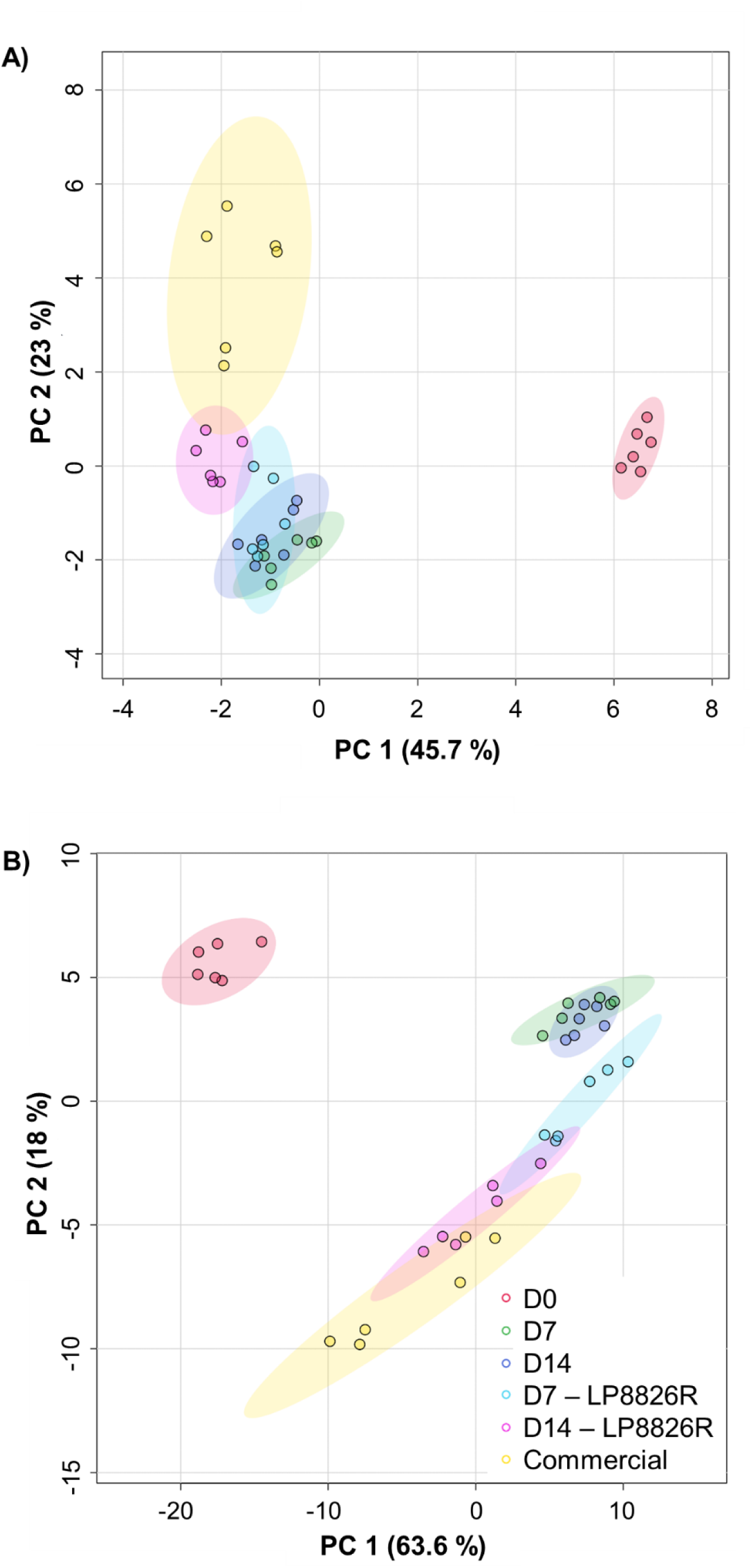
Principal component analysis (PCA) of metabolites identified by (A) GC-TOF/MS (147 metabolites) and (B) RP-LC-HRMS/MS (333 metabolites). Six replicates were tested for each condition. D0: cabbage and brine from day 0; D7 and D7 – LP8826R: laboratory scale ferments at day 7 without and with LP8826R, respectively; D14 and D14 – LP8826R: laboratory scale ferments at day 14 without and with LP8826R, respectively.

A total of 59 metabolites identified using GC-TOF/MS were significantly increased in at least one group of ferments compared to the raw cabbage homogenates (Wilcoxon rank-sum test, *p* < 0.05 and fold change ≥ 2) (**Supplemental File 4**). These compounds mainly encompassed end-products of fermentation metabolism (e.g., lactate, butane-2,3-diol), sugar and sugar alcohols (e.g., sorbitol, mannitol, trehalose, isomaltose), several amino acids (e.g., phenylalanine, tyrosine, leucine) and derivatives (e.g., PLA, ILA), and nucleobases (e.g., uracil, cytosine).

Conversely, proportions of 49 metabolites were significantly reduced in at least one group of ferments compared to the raw cabbage homogenates, including core metabolism intermediates (e.g., pyruvic acid, citric acid, alpha-ketoglutarate), sugars and sugar acids (e.g., glucose, fructose, mannose, raffinose, sucrose), and other amino acids (e.g., lysine, asparagine, glutamic acid) (Wilcoxon rank-sum test, *p* < 0.05; **Fig. 5, Fig. S4**, and **Supplemental File 4**). Based on the correlation biplots between metabolites and treatment groups performed in PCA, raw cabbage homogenates (D0) were positively correlated with the levels of sucrose, raffinose, and to a lesser extent, mannose and fructose (**Fig. S5A**). Cabbage ferments were mainly grouped together according to the proportions of fermentation end-products (e.g., sorbitol, lactic acid, mannitol, and butane-2,3-diol), PLA, ILA, and trehalose (**Fig. 5, Fig. S4**, and **Fig. S5A**). Commercial sauerkraut homogenates were further distinguished from LSF homogenates by their higher levels of putrescine and glucose-6-phosphate (**Fig. S4** and **Fig. S5A**).

**Figure 5.**
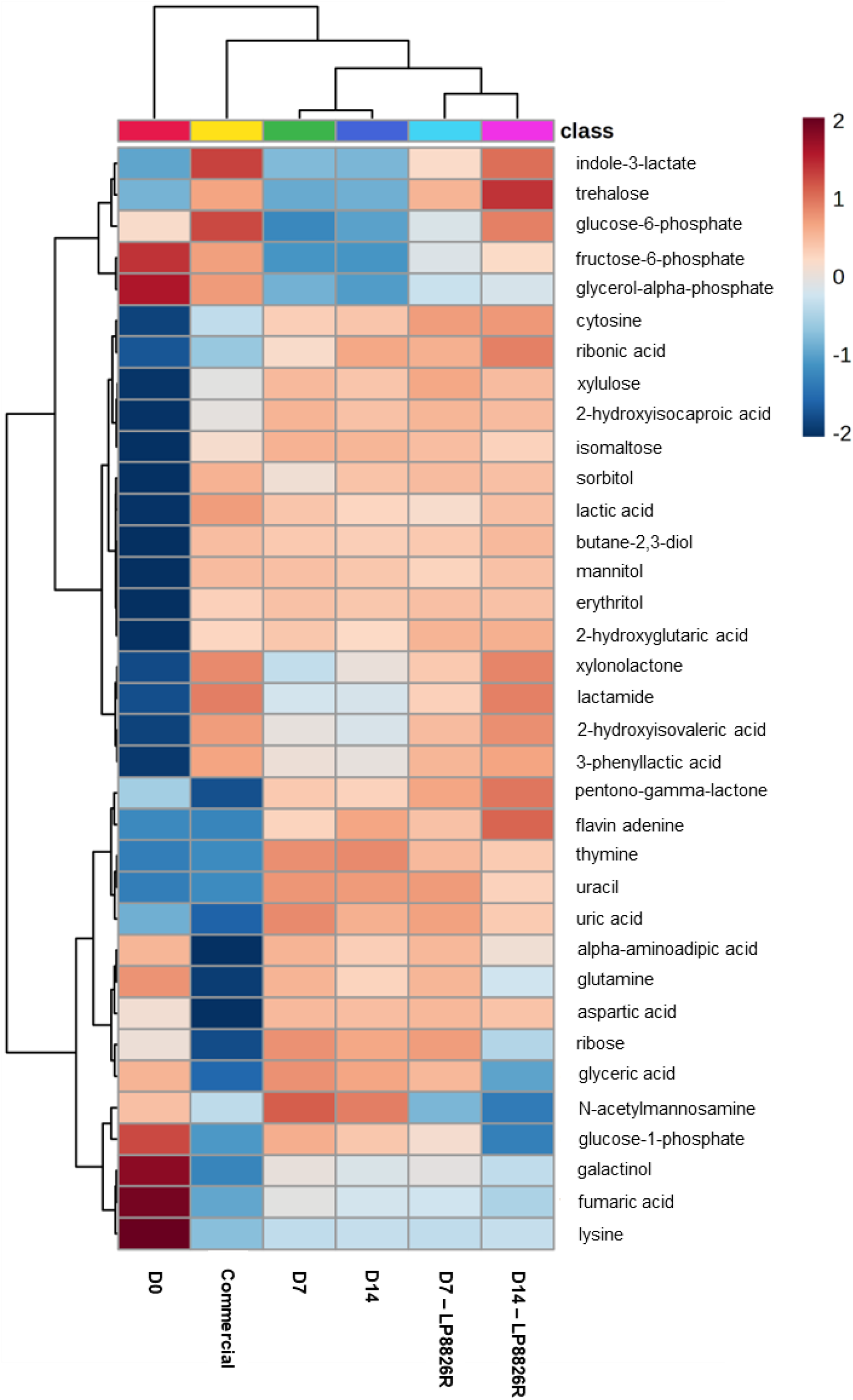
Heatmap of the relative peak heights of the top 35 metabolites detected by GC- TOF/MS that distinguish between different (fermented) cabbage homogenates. Visualization of the compounds were created using normalized data. The blue color represents lower relative peak heights while red color represents higher relative peak heights. Metabolite features were standardized through autoscaling (z-score = subtract mean and divide by standard deviation). Similarity between data points was assessed using Euclidean distance measure and then hierarchically clustered using Ward’s method. Top 35 features were determined by one-way ANOVA with Tukey’s multiple comparisons test.

RP-LC-HRMS/MS detected 6,778 metabolites in (fermented) cabbage homogenates, among which 333 metabolites were identified. As found with GC-TOF/MS, the composition and proportions (peak heights) of those identified were significantly different between the (fermented) cabbage homogenate types (PERMANOVA, *p* = 0.001, R_2_ = 0.80297) (**Fig. 4B**). The raw cabbage and brine (D0) homogenates clustered separately from fermented cabbage homogenates (*p_adj_* ≤ 0.06, pairwise Adonis test) (**Fig. 4B**). The composition and abundance of these metabolites in the LSF were similar across time points. The commercial sauerkraut was distinct from LSF (*p_adj_* < 0.05, pairwise Adonis test), except for the ferments with added LP8826R at day 14 (*p_adj_*= 0.075, pairwise Adonis test) (**Fig. 4B**). Hence, as found for GC-TOF/MS, the RP-LC-HRMS/MS metabolite profiles show that fermentation changes the metabolomes and the addition of LP8826R results in a metabolome like that found in the commercial sauerkraut (**Fig. 4B**).

While approximately one-third of the metabolites identified by GC-TOF/MS were also identified by RP-LC-HRMS/MS (e.g., amino acids, sugars, amines), this subset only constituted approximately 12% of the metabolites identified by the latter (RP-LC-HRMS/MS) technique. The other remaining metabolites mainly consisted of di-peptides, fatty acids, organo-sulfur, and poly-cyclic (e.g., epigallocatechin) compounds. In total, 254 compounds detected and identified by RP-LC-HRMS/MS were significantly increased in at least one of the fermented cabbage homogenates compared to the raw cabbage homogenates (D0) (Wilcoxon rank-sum test, *p* < 0.05 and fold change ≥ 2) (**Supplemental File 5**). Among these compounds, those also detected by GC-TOF/MS included certain amino acids (e.g., tryptophan, tyrosine) and derivatives (e.g., PLA, ILA, 2-hydroxyisocaproic acid (HICA)) and sugar alcohols (e.g., mannitol, sorbitol). The remaining compounds only detected by RP-LC-HRMS/MS mainly consisted of small di-and tri-peptides (144 out of 254 compounds; 56.7%) (e.g., phe-phe), phenolic and heterocyclic compounds (e.g., gentisic acid, 3,4-dihydroxyhydrocinnamic acid (DHCA)), other amino acid derivatives (e.g., 4-hydroxyphenyllactic acid (4-HPLA)), fatty acids (e.g., mevalonic acid), and vitamins (e.g., vitamin C, vitamin B5) (**Fig. 6**, **Fig. S6**, and **Supplemental File 5**). On the other hand, the relative abundances of 49 compounds in at least one of the ferments were significantly lower than found in the raw cabbage homogenates. Among these compounds, those also detected by GC-TOF/MS included sugars (e.g., fructose, sucrose), pyrimidines and purines (e.g., uracil, uridine), and organic acids (citric acid, gluconic acid). The remaining compounds only detected by RP-LC-HRMS/MS encompassed other sugars (e.g., turanose) and other fatty acids (e.g., (9Z)- 5,8,11-trihydroxyoctadec-9-enoic acid) (Wilcoxon rank-sum test, *p* < 0.05; **Supplemental File 5**). Based on correlation biplots between metabolites and treatment groups, the raw cabbage homogenates were distinguished from ferments by the lower relative abundance of epigallocatechin (**Fig. S5B** and **Fig. 6**). Commercial sauerkraut and LSF + LP8826R homogenates were further distinguished from other LSF homogenates by their higher relative levels of 12,13-dihydroxy-9Z-octadecenoic acid (12,13-DiHOME), 3,4-dihydroxyhydrocinnamic acid (DHCA), ILA, and PLA (**Fig. 6**, **Fig. S5B**, and **Fig. S6**). Compared to other cabbage ferments and raw cabbage homogenate, commercial ferments had higher levels of tyramine and phenylacetaldehyde (**Fig. 6** and **Fig. S6**).

**Figure 6.**
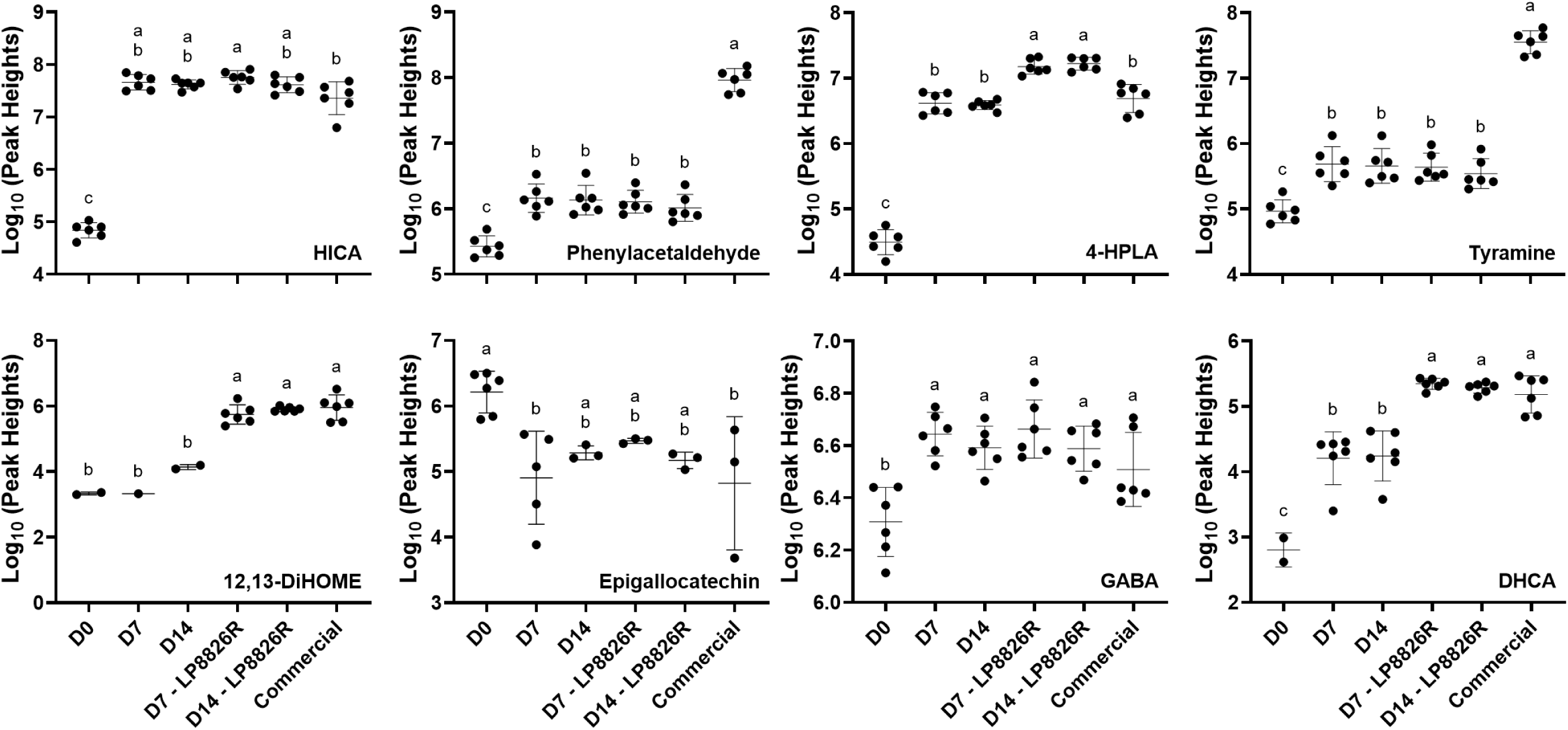
Representative metabolites detected by RP-LC-HRMS/MS. Metabolites were detected in homogenates of cabbage and brine from day 0 (D0), day 7 laboratory scale ferments (LSFs) without (D7) and with LP8826R (D7 – LP8826R), day 14 LSFs without (D14) and with LP8826R (D14 – LP8826R), and commercial ferments. Mean values not sharing letter are significantly different based on one-way ANOVA with Tukey’s multiple comparisons test. Mean ± SD for n = 6 replicates. Missing data points indicate metabolite not detected. HICA: 2-hydroxyisocaproic acid. 4-HPLA: 4-hydroxyphenyllactic acid. 12,13-DiHOME: 12,13-dihydroxy-9Z- octadecenoic acid. GABA: γ-aminobutyric acid. DHCA: 3,4-hydroxylhydrocinnamic acid.

To verify the findings of untargeted metabolomics and gain estimates of total quantities of known bioactive compounds in the ferments, targeted GC-TOF/MS was applied to measure quantities of ILA, D-PLA, and GABA, compounds found using untargeted GC-TOF/MS (ILA, D-PLA) and RP-LC-HRMS/MS (ILA, D-PLA, and GABA) (**Fig. 7**). The findings were similar to those reached by the untargeted analyses (**Fig. 5** and **Fig. 6**). Of those three compounds, GABA was detected in the highest concentrations in the raw cabbage homogenate (D0, 167.1 ± 30.4 µg/ml) but then more than doubled after fermentation (*p* < 0.0001). There were no differences in GABA quantities between the LSF and commercial ferments (**Fig. 6** and **Fig. 7**). Fermentation also resulted in significant increases in PLA for all ferments compared to the controls (D0), however, the commercial product and LSF containing LP8826R reached approximately 2-fold higher quantities (**Fig. 7**). Specifically, PLA levels increased from approximately 6 µg/ml (∼36 µM) in raw cabbage and brine (D0) to the highest concentrations of approximately 20 µg/ml (120 µM) in the LSF with LP8826R and the commercial ferments. ILA levels were significantly increased in the commercial sauerkraut and LSF containing LP8826R, but not LSF (**Fig. 7**). Particularly, ILA concentrations increased from approximately 8.5 µg/ml (∼41 µM) in cabbage and brine to the highest concentrations of around 10 µg/ml (50 µM) in the day 14 LSF + LP8826R homogenate and the commercial ferments.

**Figure 7.**
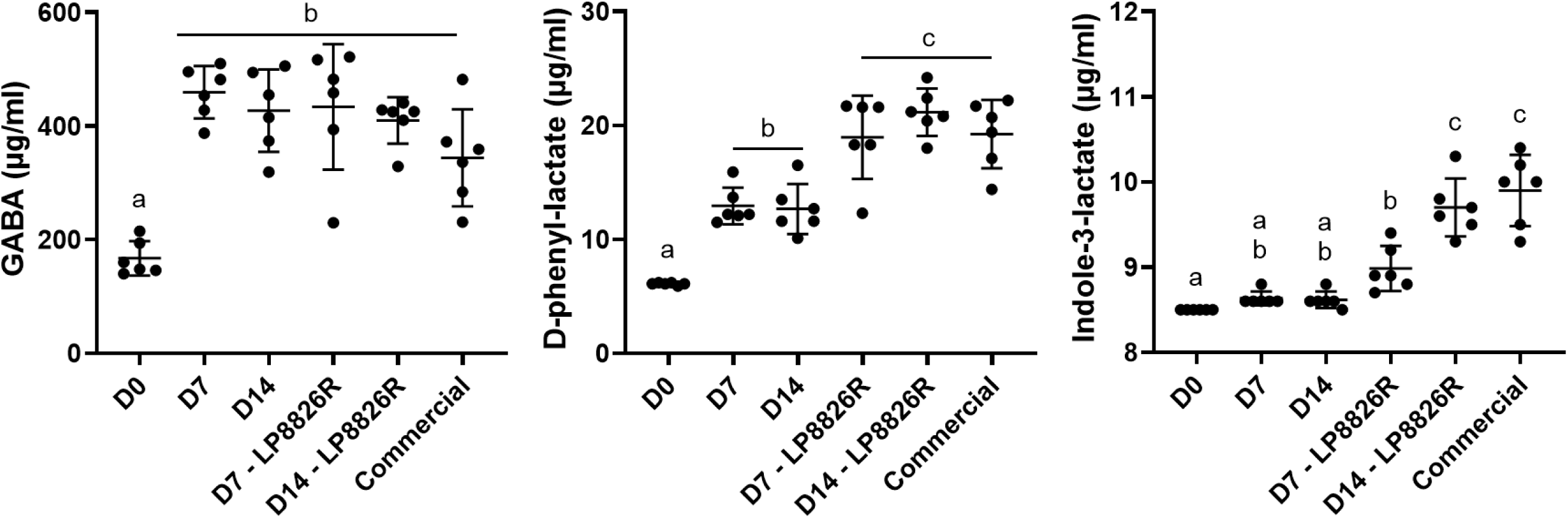
Quantities of ILA, D-PLA, and GABA in (fermented) cabbage. Mean ± SD for n = 6 replicates per condition. D0: cabbage and brine from day 0; D7/D7 – LP8826R: laboratory scale ferments with and without LP8826R from day 7; D14/D14 – LP8826R: laboratory scale ferments with or without LP8826R from day 14. Mean values not sharing letter are significantly different based on one-way ANOVA with Tukey’s multiple comparisons test.

### D-PLA, ILA, and lactate protect Caco-2 cell monolayers from cytokine-induced damage

To investigate whether individual metabolites enriched in fermented cabbage are sufficient to protect Caco-2 cell monolayers against IFN-γ and TNF-α induced damage, the monolayers were exposed to apical applications of 50 mM lactate (approximately 4.5 mg/ml), 60 µM D-PLA (approximately 10 µg/ml), or 25 µM ILA (approximately 5 µg/ml), and a mixture of all three compounds at the specified concentrations (**Fig. 8**). Metabolite concentrations used were based on the percentage of (fermented) cabbage homogenate applied onto the Caco-2 monolayers, estimates of PLA and ILA concentrations based on targeted GC-TOF/MS analysis (**Fig. 7**), and lactate concentrations found in prior studies (Gaudioso et al., 2022; Yang, Hu, Xiu, Jiang, Yang, Sarengaowa, et al., 2020). Apical treatment with these metabolites individually and combined alleviated cytokine-induced TER reductions when measured 32 h after TNF-α addition (*p* < 0.01, two-way ANOVA with Tukey’s multiple comparisons test) (**Fig. S7**). However, this beneficial effect on TER was not observed at the other time points (**Fig. 8A** and **Fig. S7**). Notably, paracellular permeability was significantly protected by the three metabolites such that the levels of FITC-dextran were significantly reduced (*p* ≤ 0.002, one-way ANOVA with Tukey’s multiple comparisons test) and comparable to the untreated controls not exposed to IFN- γ or TNF-α (*p* > 0.4997) (**Fig. 8B**). Notably, as found for the fermented cabbage, IL-8 levels were significantly elevated for Caco-2 cells exposed to the three compounds, separately and combined (**Fig. 9**). These results show the capacity of the individual metabolites to protect Caco-2 cell monolayers and that combining them did not result in a greater effect.

**Figure 8.**
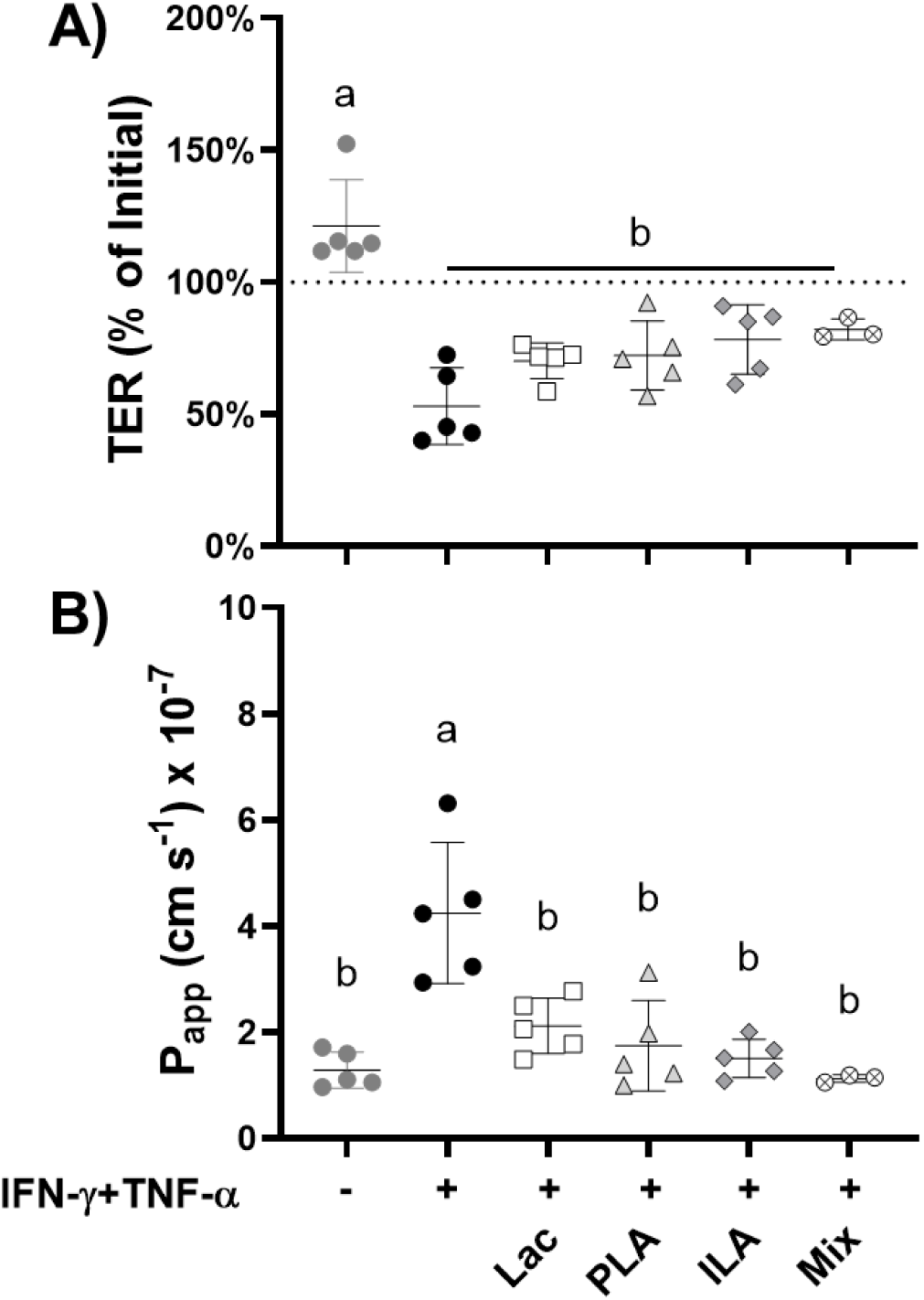
Effect of lactate, PLA, and ILA separately and combined (mix) on intestinal barrier permeability of cytokine-perturbed Caco-2 monolayers. (**A**) Trans-epithelial electrical resistance (TER) of Caco-2 monolayers 48 h after IFN-γ and TNF-α sequential additions. TER values were normalized to TER measured immediately prior to TNF-α addition. (**B**) Apparent permeability coefficient (P_app_) to FITC-dextran (4kD) of Caco-2 monolayers measured at the 48 h time point. Replicates included Caco-2 monolayers not exposed to IFN-γ and TNF-α (n = 5), controls to which the cytokines were applied (n = 5), and those exposed to the cytokines and either 50 mM lactate (Lac) (n = 5), 60 µM D-phenyl-lactate (PLA) (n = 5), 25 µM indole-3-lactate (ILA) (n = 5), or a mixture of the three metabolites (Mix) (n = 3). Mean ± SD. Mean values not sharing any letter are significantly different based on (TER) two-way or (FITC) one-way ANOVA with Tukey’s multiple comparisons test.

**Figure 9.**
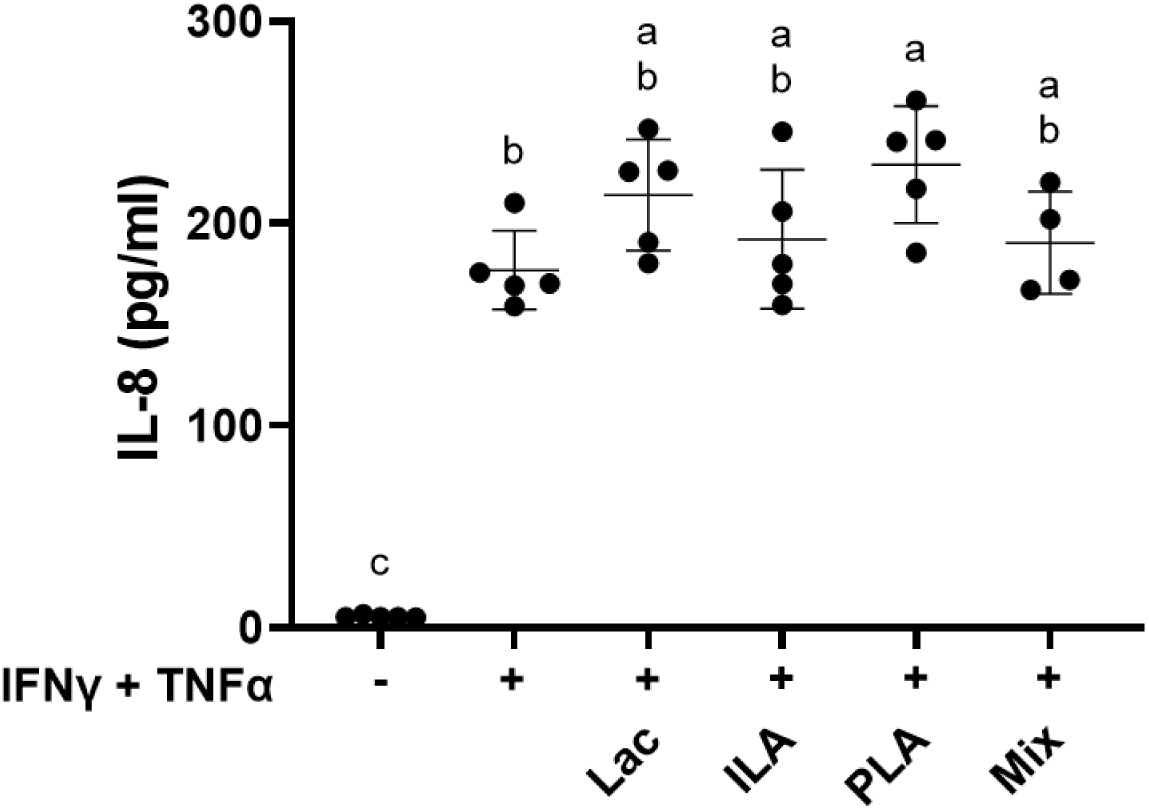
IL-8 production by Caco-2 monolayers after sequential exposure to IFN-γ and TNF-α. DMEM was collected from the basolateral side of Caco-2 monolayers 48 h after sequential application of IFN-γ and TNF-α. Replicates included Caco-2 monolayers not exposed to IFN-γ and TNF-α (n = 5), controls to which the cytokines were applied (n = 5), and those exposed to the cytokines and either 50 mM lactate (Lac) (n = 5), 60 µM D-phenyl-lactate (PLA) (n = 5), 25 µM indole-3-lactate (ILA) (n = 5), or a mixture of the three metabolites (Mix) (n = 4). The mean ± SD is shown. Letters indicate significant differences based on one-way ANOVA with Tukey’s multiple comparisons test.

## Discussion

To investigate whether fermented cabbage has the potential to support intestinal barrier function, we examined both laboratory scale and commercial cabbage ferments in a polarized Caco-2 cell model. This showed that soluble filtrates of fermented cabbage homogenates, and not raw cabbage or brine, protected against IFN-γ + TNF-α induced barrier disruption. The differences in barrier protection may be explained by the distinct metabolome profiles and enrichment of gut barrier protective compounds during fermentation. Intestinal barrier protection was retained according to FITC-dextran measurements with the application of selected fermentation-derived metabolites, D-PLA, ILA, and lactate, although they were not as effective when TER was taken into account. These findings indicate that barrier protective bioactivity of fermented cabbage likely involves the synergistic effect of multiple metabolites produced or modified by microorganisms during fermentation. These compounds were present across the ferments and were sufficient despite differences between the fermented cabbage metabolomes. This suggests that fermented cabbage contains a core metabolome that protects the intestinal barrier.

Application of a sterile filtrate of soluble fermented cabbage metabolites alleviated the increased barrier permeability induced by sequential exposure to IFN-γ and TNF-α. Fermented cabbage homogenates ameliorated cytokine-induced reductions in TER by approximately 40% by 48 h after basolateral TNF-α application and prevented cytokine-induced paracellular FITC-dextran permeability. A similar trend was also observed for filtrates of milk fermented by *Lacticaseibacillus paracasei* BL23, which resulted in a recovery of approximately 20% in TER in treated monolayers and lower permeability to FITC-dextran compared to cytokine-challenged monolayers (Zhai et al., 2019). Importantly, the addition of either 1 or 10% v/v onto the Caco-2 cell monolayers did not have a cytotoxic effect. Similarly, a shorter 24-h incubation with sauerkraut brine (10% v/v) from two artisanal producers also did not negatively affect barrier integrity, as quantified by TER (Gaudioso et al., 2022).

Despite the maintenance of barrier integrity with the fermented cabbage homogenates, there were little to no reductions in IL-8 production, indicating that the barrier protective effect of the cabbage ferments did not seem to drastically affect the inflammatory response. One notable difference with this study compared to others (e.g., Zhai et al., 2019) was that IFN-γ and TNF-α resulted in much higher (68-fold) levels of IL-8 compared to monolayers not exposed to cytokines, which may be attributed to approximately 10-fold lower baseline IL-8 level in the untreated monolayers. The approximately 1.5-fold reductions in IL-8 between fermented cabbage treated monolayers and the cytokine controls found here was comparable to the 2- to 4- fold decrease in application of either fermented milk (Zhai et al., 2019) or fermented purslane juice (Di Cagno et al., 2019). This discrepancy of barrier protection despite the increased inflammatory response may reflect signaling at different stages of NF-ĸB pathways. AHR activation, such as by LAB fermentation-derived ILA (Xia et al., 2023), and interaction with RelB may contribute to increased IL-8 levels (Vogel et al., 2007), while the NF-ĸB induced MLCK elevation may be inhibited by PPAR-γ activation (Su et al., 1999; Zhao et al., 2018) through LAB fermentation-derived ligands like PLA (Ilavenil et al., 2015). Thus, the diversity of fermentation-derived metabolites present in the cabbage homogenates may contribute to this separation between TNF-α induced NF-ĸB activation of inflammatory response (e.g., IL-8 production) and maintenance of intestinal barrier function. Future studies should focus on activation status and transcript levels of NF-ĸB subunits and other potential signaling factors (e.g., AHR, PPAR-γ) to uncover the underlying mechanisms.

Consistent with observed IL-8 levels, *TNFR2* and *CLDN2* transcript levels were significantly increased in the Caco-2 monolayers exposed with IFN-γ and TNF-α. Surprisingly, however, *MLCK* transcript levels were significantly reduced. TNF-α can increase *MLCK* transcript quantities in Caco-2 cells up to 4 h after TNF-α addition (Ma et al., 2005). Because MLCK protein levels may be sustained over longer timescales (Ma et al., 2005; Wang et al., 2005), future studies take this into consideration and should focus on quantification of protein levels following IFN-γ and TNF-α exposure.

GC-TOF/MS and RP-LC-HRMS/MS metabolomics showed that fermentation resulted in significant changes to the cabbage metabolome. Compared to the raw cabbage (D0), fermented cabbage homogenates had elevated levels of fermentation end products (e.g., lactic acid, butane-2,3-diol), organic acids (e.g., 4-hydroxyphenyllactic acid (4-HPLA)), sugar alcohols (e.g., sorbitol, mannitol, erythritol), amino acid derivatives (e.g., PLA, HICA), and fatty acids (e.g., mevalonic acid, 12,13-dihydroxy-9Z-octadecenoic acid (12,13-DiHOME)). Conversely, levels of sugars (e.g., sucrose, mannose, fructose) and metabolites involved in fermentative metabolism (e.g., citric acid, malic acid, fumaric acid, glucose-6-phosphate) were depleted. These trends during cabbage fermentation were also observed for artisanal sauerkrauts (Gaudioso et al., 2022), home fermented (Yang, Hu, Xiu, Jiang, Yang, Saren, et al., 2020), and laboratory scale Chinese northeast sauerkrauts (Hu et al., 2021; Yang, Hu, Xiu, Jiang, Yang, Sarengaowa, et al., 2020). The relative abundances of tyrosine, phenylalanine, glycine, and leucine were significantly increased after fermentation in LSFs in our study, and this was noted in artisanal sauerkraut (Gaudioso et al., 2022) and in Chinese northeast sauerkrauts, which, different than found here, also contained increases in other amino acids (e.g., aspartate, glutamate) (Yang, Hu, Xiu, Jiang, Yang, Saren, et al., 2020).

In addition to metabolites that were consistently modified across the ferments, there were compounds that were significantly altered in a limited number or individual sample types. Biplot analysis showed that LSF + LP8826R collected at day 14 were associated with higher levels of lactic acid, butane-2,3-diol, sorbitol, and mannitol. Previously, it was found that the relative abundance of *Lactiplantibacillus* in artisanal sauerkrauts correlated with the concentrations of lactic acid, acetic acid, mannitol, as well as phenylalanine and tyrosine (Gaudioso et al., 2022), highlighting that although there were overlaps, different metabolites were also enriched. As another example, the levels of putrescine and tyramine biogenic amines were significantly elevated in the commercial product. The higher levels of putrescine in commercial sauerkraut may have been partially due to storage of the product at 4 °C, since lower temperature conditions were found to be associated with increases in putrescine in sauerkraut (Peñas et al., 2010).

Targeted GC-TOF/MS quantification confirmed the presence of GABA, PLA, and ILA in the (fermented) cabbage homogenates. GABA, an inhibitory neurotransmitter, is of growing interest due to its association with neuroprotective (Hepsomali et al., 2020), anti-hypertensive (Inoue et al., 2003), and other beneficial (Hou et al., 2023; Auteri et al., 2015) properties. GABA levels reached near or above 400 µg/ml in all ferments, which is higher than detected in previous studies on fermented cabbage (Gaudioso et al., 2022; Yang, Hu, Xiu, Jiang, Yang, Saren, et al., 2020). Unlike GABA which was significantly enriched in all ferments, levels of PLA and ILA differed between the ferment sampling time and product origin. Quantities of PLA detected in the LSFs and the commercial product were comparable to levels in other vegetable ferments (e.g., sauerkraut, fermented carrots, kimchi) and higher than found some dairy ferments (Kasperek et al., 2024). Quantities of ILA detected in the LSFs and the commercial ferment were higher than those measured in other studies, including a fermented mixed vegetable juice containing cabbage, broccoli, carrot, and beetroot (Lee et al., 2021) and other vegetable (e.g., fermented carrots, fermented beets, kimchi) and dairy (e.g., kefir) ferments (Kasperek et al., 2024).

Treatment of the Caco-2 monolayers with lactate, ILA, and PLA metabolites individually and as a mixture protected against cytokine-induced increase in paracellular permeability, as measured by FITC translocation, but did not sustain protection against TER reductions 48 h after TNF-α exposure. None of the individual metabolites was able to fully recapitulate the effect of the whole cabbage ferments (10% v/v). As observed with cabbage homogenates, there were no reductions in the basolateral IL-8 quantities from monolayers treated with the individual metabolites and the mixture. Interestingly, monolayers treated with 60 µM PLA had the highest level of IL-8 production (229.0 ± 28.9 pg/ml) and monolayers treated with LSF + LP8826R collected from day 14, which had the highest PLA level (approximately 120 µM, which would estimate to 12 µM in the 10% v/v cell-free preparation) according to targeted GC-TOF/MS quantification, also had the highest IL-8 levels (150.0 ± 20.8 pg/ml) compared to other homogenates. This trend between PLA levels and IL-8 quantities were not observed for raw cabbage homogenates, which resulted in comparable IL-8 levels as LSF + LP8826R, but had the lowest PLA quantities. It is notable that the metabolite concentrations applied to the Caco-2 monolayers were higher than physiologically relevant levels that were shown to activate the cellular receptor for these metabolites. Specifically, the half maximal effective concentrations (EC_50_) for hydroxycarboxylic acid receptor 3 (HCA3) activation by PLA and ILA are in the 0.1 to 5 µM range (Peters et al., 2019) and lactate activation of the HCA1/GPR81 receptor in the 5 to 20 mM range (Koh et al., 2016). Moreover, cell culture and animal model studies reported intestinal barrier protective properties following administration or production of microbiota-associated PLA (Shelton et al., 2023; Kim et al., 2019), ILA (Henrick et al., 2021; Zhang et al., 2023; Xia et al., 2023), and lactate (Iraporda et al., 2015; Lee et al., 2018; Li et al., 2024; Yu et al., 2021). These effects were noted to be modulated through activation of signaling pathways involving PPAR-γ (PLA; (Shelton et al., 2023)), AHR (ILA; (Cui et al., 2023)), and GPR81 (lactate; (Li et al., 2024)). Thus, at least for protection against IFN-γ and TNF-α as measured in the Caco-2 monolayer model here, the complexity of metabolites in the whole food (filtered homogenate) was more protective compared to the individual metabolites.

Notably, LP8826R addition affected the LSF cabbage metabolomes and Caco-2 responses such that those ferments better resembled the commercial product. LP8826R dominated the total culturable LAB population three days after addition, leading to elevated total LAB numbers and lower brine pH compared to the LSF without LP8826R. A similar result was found upon the addition of a *L. plantarum* strain to Chinese northeastern sauerkraut fermentations (Yang, Hu, Xiu, Jiang, Yang, Sarengaowa, et al., 2020). These findings show the potential application of adding probiotic starter cultures in vegetable fermentations (Patarata et al., 2024). Furthermore, remarkably, despite the differences in metabolomes among the cabbage ferments, all cabbage ferments exhibited intestinal barrier protective property, indicating the presence of a fermentation-derived core metabolome that confer bioactivity.

Based on these findings, forthcoming studies should elucidate the specific metabolites and signaling pathways via which fermented cabbage affects intestinal epithelial cell function and assess the impacts when the ferments with viable microorganisms present. Verifying the potential health-promoting properties of fermented fruit and vegetable foods can also support the identification of starter culture strains which can be added to the ferment to confer desired bioactive activities. Ultimately, human studies are required to determine the intake recommendation of fermented cabbage that will yield physiologically relevant impact in the target populations.

## Author Contributions

**Lei Wei** contributed to the preparation of the original draft, reviewing and editing, conceptualization, and visualization. **Maria L. Marco** contributed to reviewing and editing of the draft, conceptualization, project administration, supervision, and funding acquisition.

## Data Availability

The GC-TOF/MS and RP-LC-HRMS/MS peak height datasets used in the untargeted metabolomic analysis are available in the Harvard Dataverse repository at https://doi.org/10.7910/DVN/EM6HXV.

## Acknowledgements

This work was funded by Specialty Crop Block Grant from California Department of Food and Agriculture (CDFA) (19-0001-050-SF) and the Jastro and Shields Graduate Research Award from the University of California, Davis. We would like to thank the UC Davis West Coast Metabolomics Center for performing the (un)targeted metabolomics and providing feedback on the metabolomics methods section.

